# Feature-Mask-Based Strategies for Subtype-Specific Freezing of Gait Detection using CNNs

**DOI:** 10.1101/2025.05.26.656145

**Authors:** Xinyue Yu, Kaylena Ehgoetz Marten, Arash Arami

## Abstract

**Objective:** Freezing of gait (FOG), a disabling symptom of Parkinson’s disease, varies in manifestations and motion contexts. Its heterogeneity motivates subtype categorization such as manifestation-specific subtypes (akinesia, trembling or shuffling) or motion-specific subtypes (gait-initiation, walking or turning). Despite numerous promising deep learning FOG detection studies, few consider FOG heterogeneity. It remains unclear whether different subtypes require distinct detection strategies, and whether tailoring subtype-specific models could enhance detection generalizability across subtypes. Methods: To address these questions, we categorize FOG data into manifestation- or motion-specific subtypes and derive their corresponding detection strategies as interpretable feature masks. We then propose a feature-mask-based CNN that explicitly embeds the identified strategies. Using waist-mounted 3D accelerometer data, a general CNN and subtype-specific CNNs are trained. Results: According to feature-mask analysis, motion-specific subtypes share a common detection strategy, whereas manifestation-specific subtypes require distinct strategies. Manifestation models exhibit enhanced generalizability across subtypes compared to the general model, boosting the overall average FOG detection sensitivity by 24.95%±9.80% and specificity by 18.29%±8.71%. Conversely, motion models reduce the overall FOG sensitivity by 1.89%±8.74% and specificity by 5.17%±10.76%. Conclusions: The detection strategy is mainly driven by manifestation composition of the data. The general model favors the dominant manifestation-specific subtype group(s), a bias corrected by tailored manifestation-specific strategies. No comparable benefit arises from motion models due to their similar manifestation compositions. Significance: This study interpretably reveals the detection strategies required by different FOG subtypes and demonstrates the effectiveness of subtype-specific tailoring in improving FOG detection generalizability.

## I. Introduction

Parkinson’s disease (PD) is a neurodegenerative disorder causing different motor deficits such as freezing of gait (FOG) [1]. FOG, common in advanced PD patients [2], is characterized by temporary, episodic reduced ability or inability to move forward, despite the intent to walk [3]. Typical pathophysiologies of FOG exhibit disturbed temporal and frequency patterns, as evidenced by leg trembling in 4 to 5 Hz [4], impaired gait timing regulation [5], and asymmetrical and discoordinated gait [6]. Beyond its debilitating impact on quality of life, FOG is the leading cause of falls in PD patients [7].

FOG episodes vary in their responsiveness to dopaminergic medication, manifestation, and context, leading to the definition of subtypes within each aspect. Pharmacological subtypes categorize FOG as either responsive or unresponsive to levodopa [8], a cornerstone dopamine therapy for PD [1]. Based on manifestations, FOG has been categorized into gait [9] or phenomenological [10] subtypes, which include shuffling-forward, trembling in-place, and akinesia (complete lack of movement, the rarest [9]). Contextually, FOG stems from internal (e.g., affective state, cognitive load) and external factors. For instance, FOG may occur with or without heightened emotional arousal, as found in an investigation that 48% of 29 freezers reported panic attacks during FOG [11]. Similarly, FOG may occur with or without added cognitive load, as demonstrated in [12], where FOG was observed both with and without a dual-task color classification during turning. As an example of environmental-driven external factors, doorway width is inversely related to frequency of FOG episodes [13]. As an instance of interaction-driven external factors, motion context also influences FOG, with its episodes observed during gait initiation [14], walking [15], and turning [16].

Given its debilitating impact, FOG detection has been actively studied. Leveraging wearable sensors, detection algorithms can help FOG monitoring, auto-annotation of clinical assessment, and on-demand activation of nonpharmaceutical FOG treatments, such as cueing via vibration or audiovisual stimuli [17] and assistive exoskeletons [18]. Kinematics-capturing wearables, primarily inertial measurement units (IMUs) or accelerometers, are the most widely used sensors in FOG studies [19], making kinematics-based detection our focus.

One of the earliest kinematics-based FOG detection methods uses empirical thresholds of the freeze index (FI) (ratio of power in the freeze band (3–8 Hz) to the locomotor band (0.5–3 Hz) of vertical leg acceleration [20]), which is simple and widely used but may misclassify voluntary stops as FOG due to similar spectral shifts that increase FI by decreasing locomotor band power [21]. Then, machine learning models such as support vector machines [22] and decision trees [23] demonstrate enhanced detection performance by incorporating diverse hand-engineered features [24], [25]. However, feature extraction requires prior knowledge and thus may overlook some useful though less interpretable data-driven patterns. Recently, popular deep learning (DL) methods such as feedforward probabilistic neural network [25], recurrent neural network [26], and Convolutional Neural Network (CNN) [27] have been applied to FOG detection problem. They have shown impressive detection accuracy by automatically capturing complicated and various data features, yet their success depends on large, diverse training datasets [15], challenging to collect due to FOG’s episodic nature.

Despite promising results of DL methods in kinematics-based FOG detection [25] [28] [29] [30], only few approaches account for FOG heterogeneity in their design. Mo and Chan trained a transformer to classify non-FOG, FOG at walking, FOG at turning, or FOG at gait initiation [31]. Yang et al. evaluated different deep models to differentiate trembling-in-place FOG, akinesia FOG, and non-FOG using lower-body movement recorded by IMUs (where shuffling treated as non-FOG events) [32]. These studies show that FOG episodes under different motion contexts or with different manifestations are distinguishable, supporting the feasibility of subtype-specific detection. Shaban et al. trained multiple CNNs, each focusing on FOG detection at walking, turning or gait initiation, and found the turning CNN performed the worst, suggesting greater FOG pattern complexity at turning [33]. However, without a comparison to a general model, it remains unclear whether subtype-specific training offers any enhancement.

Considering FOG heterogeneity is essential for designing effective detection algorithms, as subtype importance may vary among patients. For instance, some may find akinesia the most disruptive, while others may be more concerned with trembling, the typically most frequent manifestation [9]. Without maintaining detection performance across different subtypes, detection algorithms may fail to generalize to diverse patient needs. Additionally, DL models trained on imbalanced data may be prone to bias, overfitting to common FOG subtypes while neglecting less frequent cases such as akinesia. A balanced approach that detects all subtypes reliably also offers unbiased support for future research, such as whether different FOG subtypes have different neuropathologies [10]. This study focuses on FOG heterogeneity in motion context and manifestations, since both are directly linked to kinematics. We define motion-specific subtypes as gait initiation, walking, and turning, and manifestation-specific subtypes as shuffling, trembling, and akinesia.

Given the limited investigation of FOG heterogeneity in DL-based FOG detection, we first examined whether each subtype requires a unique feature-based detection strategy. We then propose a featuremask-based CNN that explicitly and interpretably incorporate these strategies to assess whether tailored models improve detection generalizability across subtypes compared to a general model.

## II. Materials and Methods

We used tDCS FOG dataset [34], consisting of 833 trials (215.25 hours) collected from 62 PD patients performing Ziegler FOG-provoking protocol [35]. Each trial began with quite standing, followed by a 1 m walk to a test spot for two 360° in-place turns (clockwise and counter-clockwise), then a 2 m walk through an open doorway and back. This protocol was conducted in both ON and OFF dopaminergic medication states. All trials were treated collectively, as DL-based FOG assessment models can generalize across medication states [30]. The 3D acceleration of the participants’ lower back was acquired at 128 Hz along with expert-annotated labels of non-FOG (67.24%), and FOG (32.76%). The FOG samples were further annotated as FOG in turning (76.91%), walking (9.50%), or gait initiation (13.59%), corresponding to motion-specific subtypes.

Fig. 1 illustrates the methodological pipeline. After pre-processing and feature extraction, we split the dataset into either manifestation- or motion-specific subtype groups, enabling us to derive and compare feature masks that represent subtype-specific and general detection strategies. Finally, models embedding those feature masks were trained and compared to assess gains from subtype-tailored modeling.

**Fig. 1.**
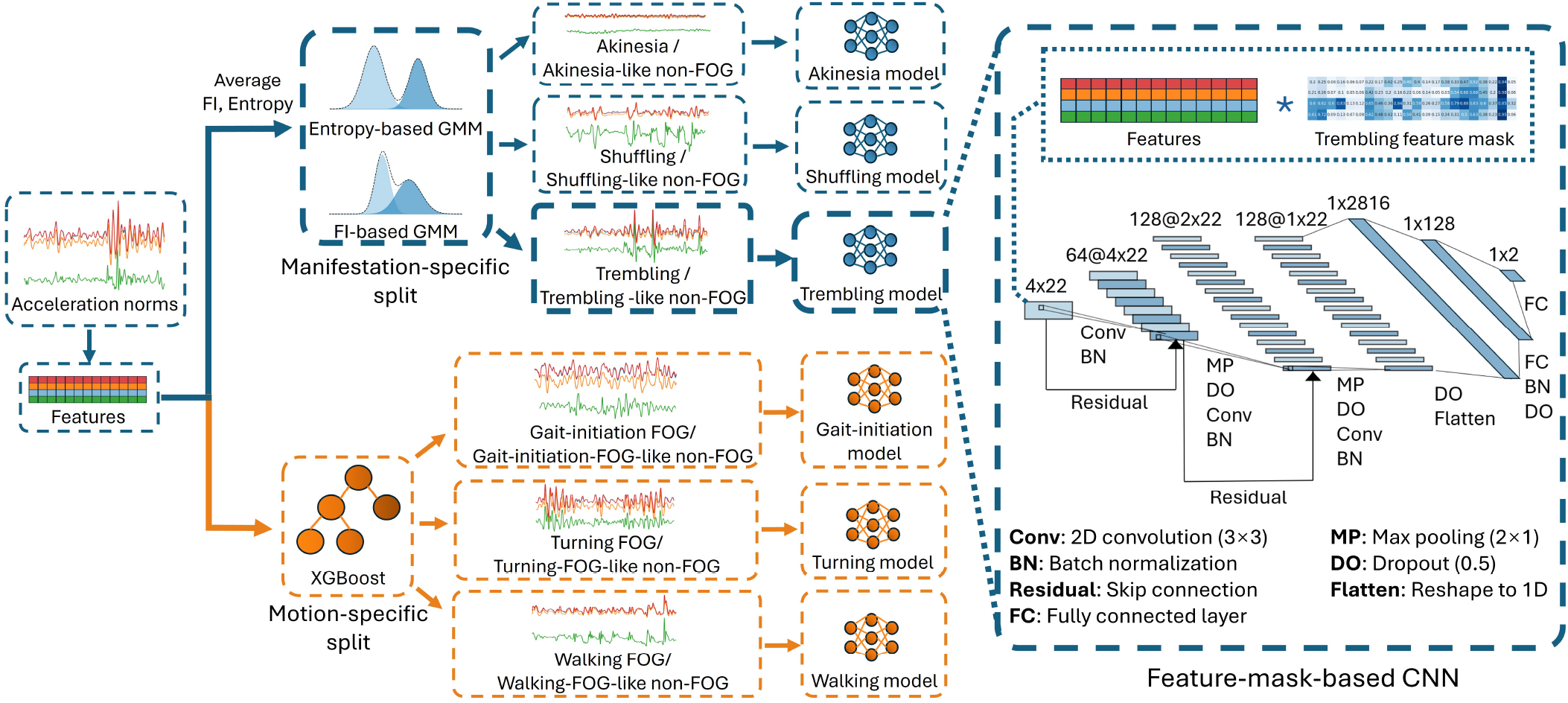
Methodological pipeline with feature-mask-based CNN architecture. For each window, acceleration norms are computed to extract a feature matrix. The window is then assigned to a manifestation-specific subtype group (akinesia, shuffling, trembling) via a GMM-based split, or a motion-specific subtype group (gait initiation, turning, walking) via an XGBoost-based split. For model training, a feature mask is generated for each subtype group and the corresponding feature-mask-based CNN is then trained with it. Each CNN takes the element-wise product of the feature mask and feature matrix as input, followed by three convolutional layers with batch normalization, residual connections, max pooling, and dropout, and concludes with fully connected layers for FOG/non-FOG classification. During model inference, the CNN (e.g. trembling model) corresponding to the subtype group (e.g. trembling) of the input window is activated.

### A. Data Preprocessing

The raw data were first segmented into 4s sliding windows with 0.25s overlap. A relatively large window size was chosen to catch gait changes while a small overlap helps capture short FOG episodes. 3D acceleration norm (*acc*_*total*_) and 2D norms in the sagittal (*acc*_*sag*_), transverse (*acc*_*trans*_), and frontal (*acc*_*front*_) planes were used to reduce sensitivity to sensor placement variations and enable more targeted analysis. Ground truth labels for non-FOG, FOG, and motion-specific subtypes were assigned to a window using Algorithm 1, a flexible labeling method designed for faster detection than traditional majority voting.

The dataset was then split into 10 trial-independent folds (patients may appear in multiple folds), using Algorithm 2, which balances sample counts of non-FOG, FOG, motion-specific subtypes, and short FOG across folds. For 10-fold cross-validation, a 80%-10%-10% train-validation-test split was achieved by each time setting 8 folds as training, one fold as validation, and one fold as test set. The roles of the folds rotated sequentially across iterations.

#### Algorithm 1

Dominant-Frequency-Based Labeling

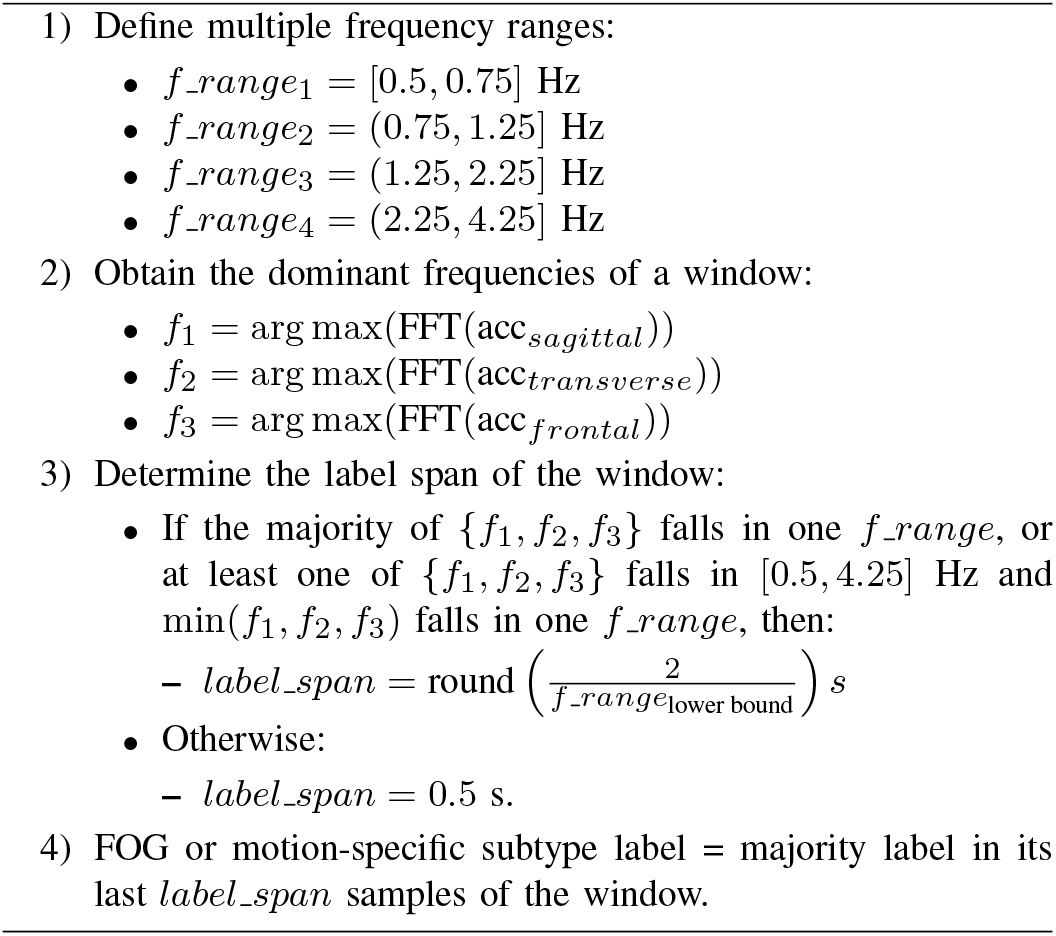

#### Algorithm 2

Balanced Fold Assignment

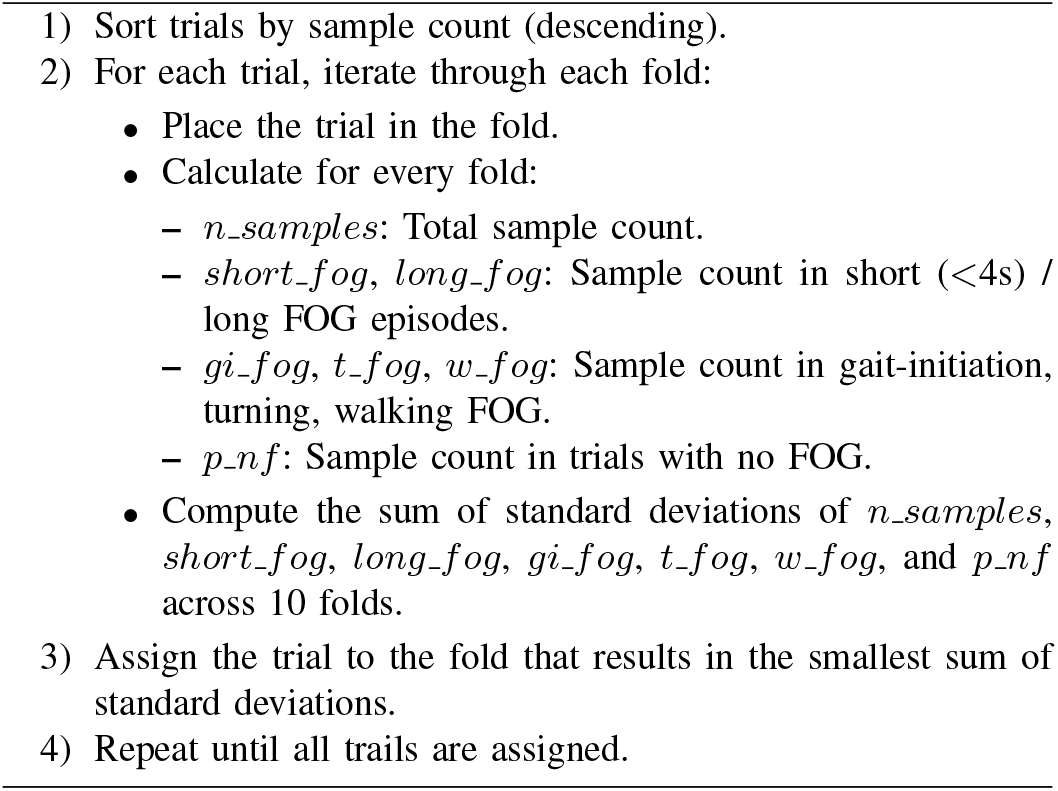

### B. Feature Extractions

16 time-domain (TD) and 6 frequency-domain (FD) features commonly used for FOG detection algorithms [36] [37] (Table I) were computed for each of the four acceleration norms of each window. Since majority of the spectral energy of human motion lies below 15 Hz [38], the FD features were extracted from the 0.5-15 Hz range of the Fast Fourier Transform (FFT) spectrum using a Hanning window.

**TABLE I.** Summary of Time-domain and Frequency-domain Features.

| Time Domain (TD) | Description |
| --- | --- |
| Mean | Average value |
| Median (Med) | Middle value of the data when sorted in ascending order |
| Trimmed median (TremMed) | Median value after removing 5% of the lowest and highest data |
| Standard deviation (Std) | Variation from the average |
| Min, Max | Minimum and maximum values |
| Range | Difference between the maximum and minimum values |
| Root mean square (RMS) | Square root of the average of the squared values |
| Mean absolute deviation (MAD) | Average of the absolute differences from the mean value |
| 25th percentile (P25), 75th percentile (P75) | The value below which 25% or 75% of the data fall into |
| Interquartile Range (IQR) | Difference between P75 and P25 |
| Skewness (Skew) | Asymmetry of the data distribution |
| Kurtosis (Kurt) | Sharpness of the peak, with high Kurtosis corresponding to more outliers. |
| Normalized Signal Magnitude Area (NormSMA) | Sum of the absolute values, normalized by window length |
| Mean Crossing Rate (MCR) | Number of times that the data crosses its average |
| Frequency Domain (FD) | Description |
| Spectral Entropy (SpecEnt) | Randomness and complexity of the frequency components of the data |
| Locomotion Band Power (LBPr) | Energy in 0.5 to 3 Hz frequency range |
| Freeze Band Power (FBPr) | Energy in 3 to 8 Hz frequency range |
| Freezing Index (FI) | Ratio of FBPr over LBPr |
| Peak Frequency (PeakFreq) | Frequency associated with the maximum power spectrum value |
| Average Energy (AvgEng) | Mean value of the energy of a window |

### C. Subtype-Driven Data Split

While the dataset includes motion-specific labels for FOG, it lacks such labels for non-FOG and manifestation-specific labels. To enable subtype-specific modeling, using train-set FOG data, data split methods were developed to assign each FOG window to a manifestation- or motion-specific subtype. The resulting split methods were then applied unchanged to non-FOG windows, aiming to assign each to the manifestation- or motion-specific subtype it most resembled and thus was most likely to be mistaken for as FOG. As a result, each subtype group contained its corresponding FOG subtype and the non-FOG samples similar to that subtype.

#### 1) Manifestation-Specific Split

Akinesia, characterized by a complete lack of movement [9], was distinguished from the other two motion-heavy subtypes using spectral entropy, which measures the complexity and orderliness of a signal. To further differentiate between trembling and shuffling, we utilized FI, which quantifies the ratio of power in shuffling-focused locomotion band (0.5 to 3 Hz) to the trembling-focused freezing band (3 to 8 Hz) [20]. Although originally defined for vertical acceleration, the extension of FI to other axes also aids FOG characterization [25]. We fitted two two-cluster-Gaussian Mixture Models (GMM), via Expectation Maximization, separately on the average spectral entropy and FI across four acceleration norms. We chose GMM for its soft probabilistic assignments which enable flexible subtype separation. Windows in the low-entropy cluster of entropy-based GMM were labeled akinesia. Among the rest, those in the low-FI cluster of FI-based GMM were labeled shuffling, and the others were labeled trembling.

#### 2) Motion-Specific Split

An Extreme Gradient Boosting (XG-Boost) classifier was trained on extracted features and dataset-provided movement labels to assign each FOG window into a motion-specific subtype. XGBoost was chosen for its lightweight implementation and interpretable feature importance, aligning with manifestation-specific split.

### D. Feature Mask and Feature-Mask-Based CNN

#### 1) Effect-Size-Based Feature Mask

To mitigate sample size effects, effect size, calculated as the mean difference between groups divided by the pooled standard deviation, was used instead of p-value to evaluate feature differences between FOG and non-FOG groups [39]. A 4 × 22 feature mask was obtained by min-max normalizing an effect size matrix, where each element corresponded to a feature in Table I. The feature mask, as later embedded into a detection model, served as an importance weight matrix highlighting features with greater differences between FOG and non-FOG groups and directly visualizing the model’s detection strategy.

All feature masks were generated using train set. For the general model, a general feature mask was obtained using all data. For each subtype model, a subtype-specific feature mask was obtained with data of the corresponding subtype group. Spearman’s rank and Pearson correlations were calculated between every pair of these feature masks to assess their similarity in feature ranking and relative feature importance, respectively. For analysis, feature masks were also generated for each FOG subtype and all non-FOG.

#### 2) Feature-Mask-Based CNN Architecture

As illustrated in Fig. 1, the model input consists of the element-wise multiplication between the extracted feature matrix and its corresponding feature mask, which is then fed into three convolutional layers to catch relations across features and across acceleration norms. Batch normalization, residual connection, and dropout layer between each convolutional layer were used to stabilize training and avoid overfitting. Flattened convolutional outputs pass through two fully connected layers, yielding the FOG and non-FOG probability.

#### 3) Model Training and Evaluation

Subtype-specific train, validation, and test sets were obtained by applying the data split methods separately to the train, validation, and test sets. The general model was trained on all data, while each subtype model was trained using its corresponding subtype-specific train and validation set. The learning rate was set at 0.001. Cross-entropy loss and Adam optimizer were used. Down-sampling was applied to the majority class on training and validation sets to address class imbalance. Early stopping was employed to prevent overfitting, with a patience (number of epochs to wait after the validation loss plateaus before halting training) of 20 for general model and 40 for subtype model, given its smaller dataset. The training and validation loss curves were also monitored to ensure both decreased smoothly without significant divergence, indicating neither underfitting nor overfitting.

Each subtype model was then evaluated on its corresponding subtype test set to reflect subtype-based model activation, while the general model was evaluated on all subtype test sets. with FOG as positive class, we reported ten-fold true positive rate (TPR), i.e., sensitivity, and true negative rate (TNR), i.e., specificity. For each manifestation and motion test sets separately, an Aligned Rank Transform Analysis of Variance (ART ANOVA) was performed on TPRs and TNRs to examine the effects of model and test set and whether within-test-set comparisons were justified. Two-sided Wilcoxon signed-rank tests with a Bonferroni correction factor of 3 were used to assess the statistical significance of TPR and TNR differences between each subtype model and the general model.

## III. Results

### A. Data Split

Across 10 folds, the silhouette score was 0.444 ± 0.004 for entropy-based GMM and 0.671 ± 0.003 for FI-based GMM, indicating respectively a weak-to-moderate and moderate inter-cluster distinctiveness. The resulting manifestation-specific split yielded a subtype distribution of 15.03% ± 1.12% akinesia, 41.32% ± 0.69% shuffling, and 43.55% ± 0.79% trembling, and a corresponding non-FOG distribution of 11.99% ± 0.53% akinesia-like, 84.33% ± 0.43% shuffling-like, and 3.68% ± 0.16% trembling-like.

Over 10 folds, RMS and TremMed of *acc*_*trans*_ are the main features used by XGBoost. XGBoost achieves a near-perfect train accuracy of 0.990±0.002, making its split on train set equivalent to using true labels. Its test accuracy (0.765±0.047) is lower, reflecting the uncertainty in model activation during practical inference. The resulting motion-specific split yielded a non-FOG distribution of 5.29% ± 0.94% gait-initiation-FOG-like, 92.09% ± 0.95% turning-FOG-like, and 2.62% ± 0.55% walking-FOG-like.

### B. Feature Mask Visualization

The average feature masks across 10 folds are visualized in Fig. 2 and Appendix I. All exhibit low variance, as indicated by a moderate or below-moderate coefficient of variation (CV) of 0.2-0.3 for all features with notable significance (normalized effect size *>* 0.1).

**Fig. 2.**
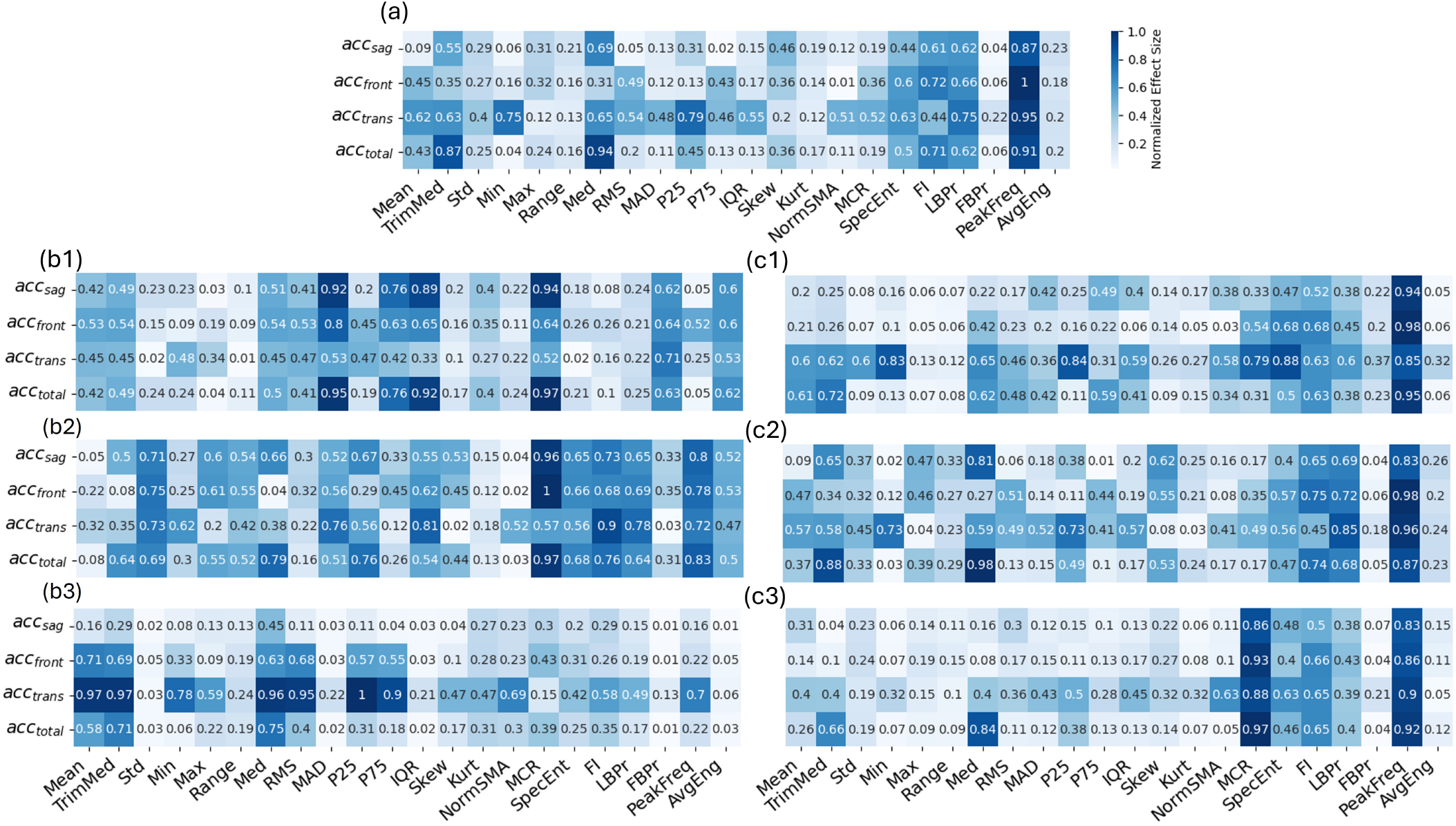
Average feature masks obtained with (a) all data, (b) manifestation-specific groups ((b1) Akinesia, (b2) Shuffling, (b3) Trembling), and (c) motion-specific groups ((c1) Gait-initiation, (c2) Turning, (c3) Walking). Manifestation-specific feature masks (b) differ notably from each other and the general mask (a), whereas motion-specific feature masks (c) resemble each other and the general mask (a).

Manifestation feature masks in Fig. 2 (b) show distinct pattern from each other and from the general feature mask in (a). Specifically, while the general feature mask attributes higher importance to FD features (e.g., PeakFreq), akinesia feature mask highly values TD features (e.g., MAD and IQR). Shuffling feature mask exhibits a more uniform feature importance distribution across TD and FD. Trembling feature mask pays even higher attention to TD features related to the transverse plane. In contrast, motion feature masks in (c1-3) share a relatively similar pattern to each other and to the general feature mask. These feature masks rely heavily on FD features, with similar ranked preferences in peak frequency, FI, locomotion-band power, and spectral entropy as the top features. The only noticeable, albeit small, difference is the higher reliance on mean crossing rate by walking feature mask. These observations hold when extending the non-FOG coverage to all non-FOG (Appendix I), which shows that the same-group non-FOG is the main non-FOG focus of the presented subtype-specific detection strategy.

### C. Feature Mask Similarity

Fig. 3 displays the 10-fold average of Spearman’s rank and Pearson correlations between feature masks. These average correlations have a low variance across 10 folds with highest CV of 0.17.

**Fig. 3.**
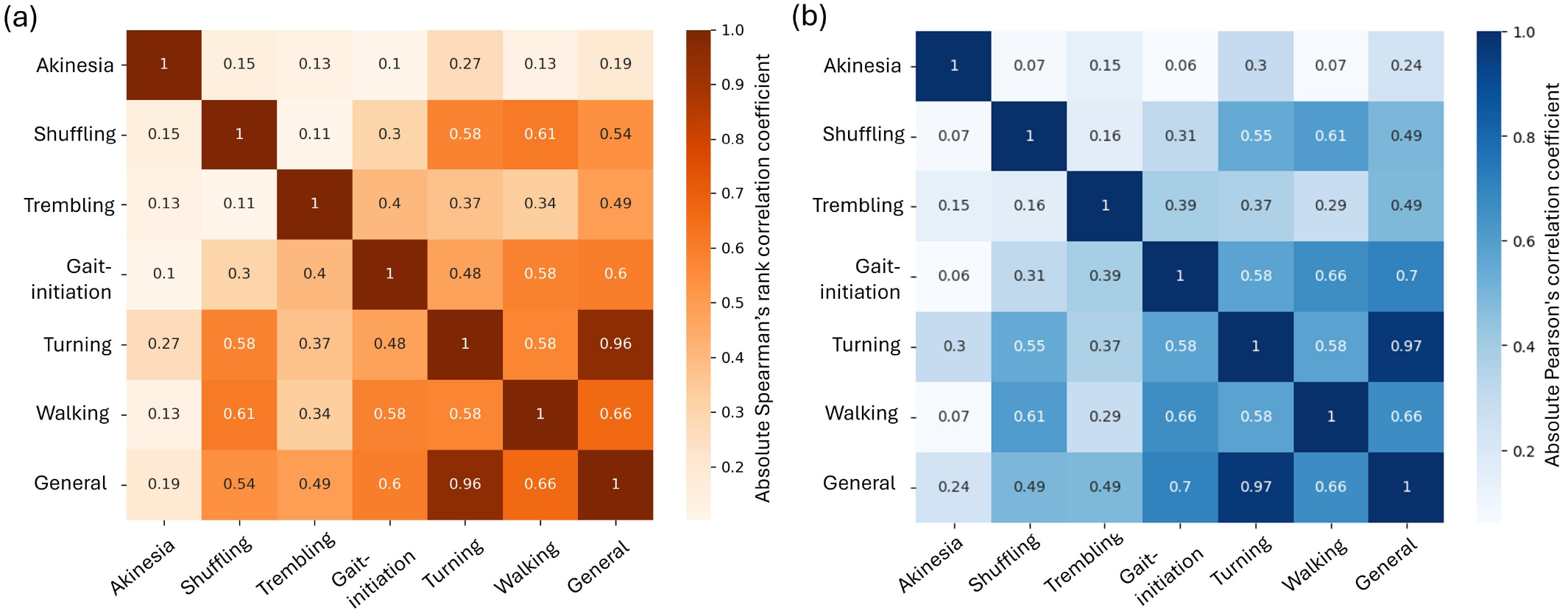
Spearman’s rank correlation (a) and Pearson correlation (b) between feature masks. It quantitatively confirms that manifestation feature masks have higher inter-subtype and compared-to-general distinctiveness, indicated by lower correlations, than motion feature masks.

Quantitatively confirming their higher distinctiveness from the general feature mask, manifestation feature masks exhibit lower correlations to the general feature mask (Spearman: 0.19 to 0.49; Pearson: 0.24 to 0.49) than motion feature masks (Spearman: 0.6 to 0.96; Pearson: 0.66 to 0.97). Among manifestation feature masks, shuffling has the highest Spearman correlation with the general feature mask. Shuffling and trembling feature masks both show the highest Pearson correlation with the general mask. Across motion feature masks, turning shows the highest Spearman and Pearson correlation with the general mask. The higher inter-subtype distinctness of manifestation feature masks is also quantitatively validated by their lower inter-subtype correlations (Spearman: 0.11 to 0.15; Pearson: 0.07 to 0.16) than those of motion feature masks (Spearman: 0.48 to 0.58; Pearson: 0.58 to 0.66),

### D. Feature-Mask-Based CNN FOG Detection Performances

Table II shows the ART ANOVA TEST results. Fig. 4 presents the 10-fold TPRs and TNRs on each subtype test set by the corresponding subtype model and the general model, while Fig. 5 illustrates the average differences between them.

**TABLE I.**
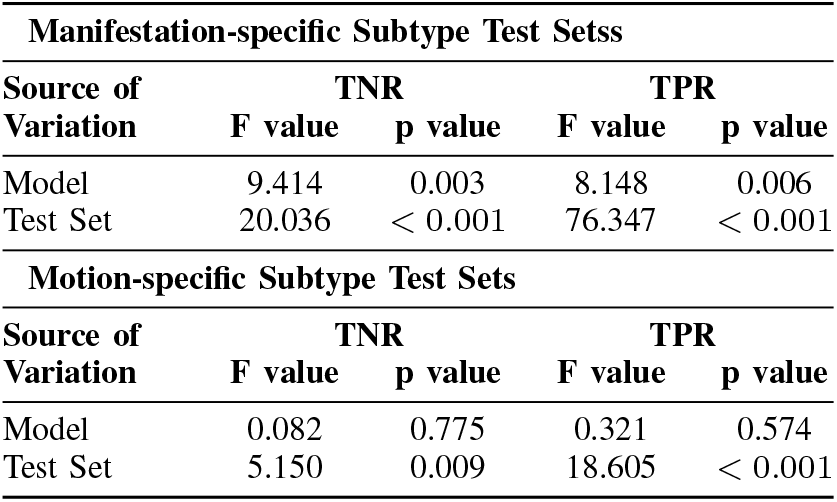
ART ANOVA Test Results.

**Fig. 4.**
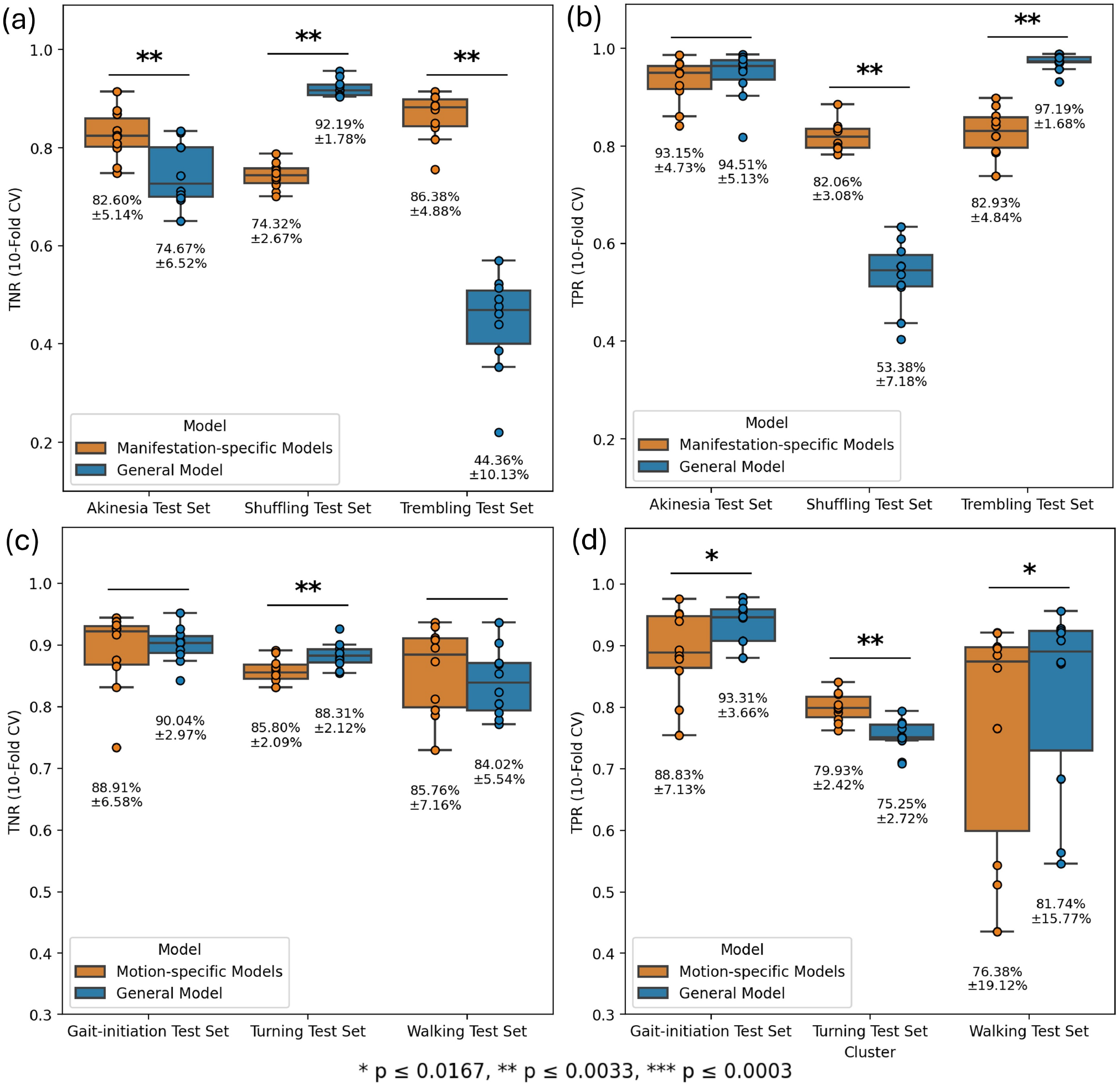
10-fold performance comparison between manifestation-specific models vs general model on (a) TNR (b) TPR, and motion-specific models vs general model on (c) TNR (d) TPR. Mean ± standard deviation is annotated. Manifestation-specific models significantly improve TPR of shuffling and TNR of akinesia- and trembling-like non-FOG. Motion-specific models modestly improve TPR of turning FOG.

**Fig. 5.**
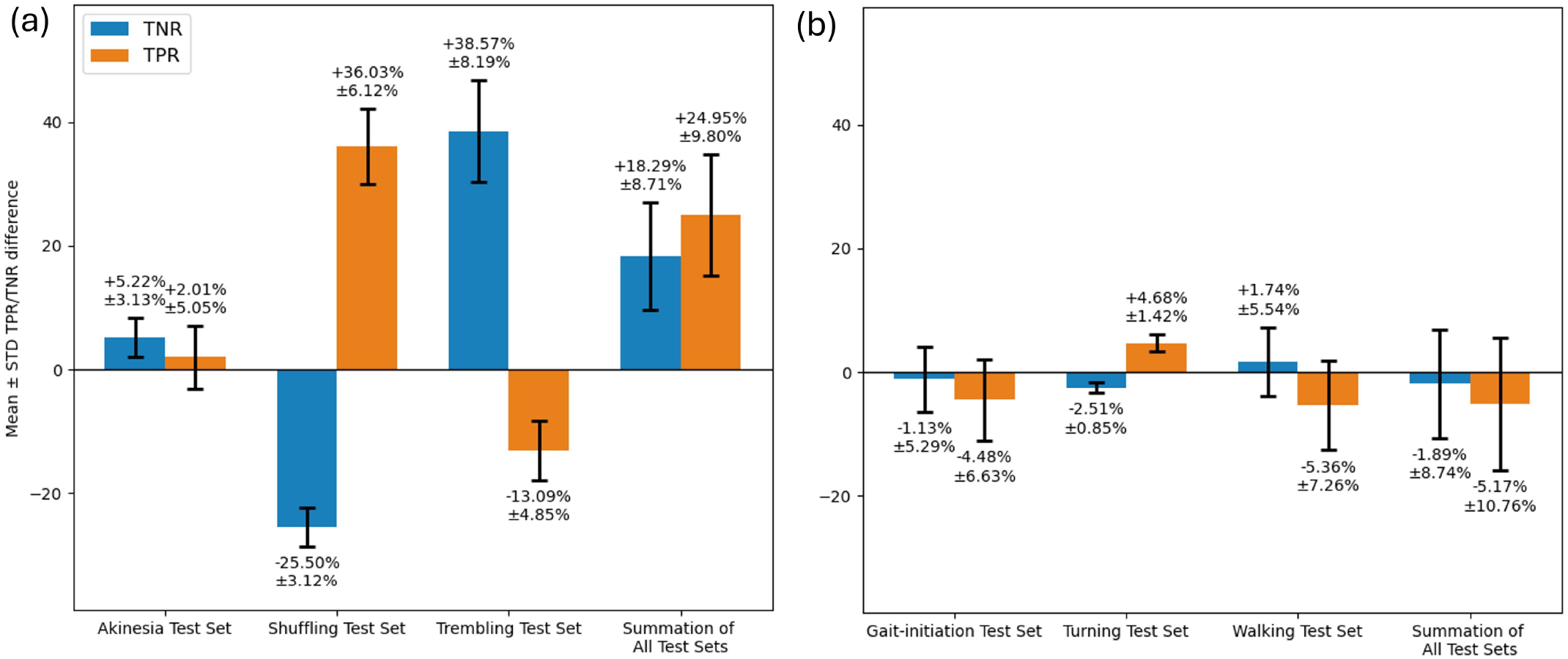
10-fold average TPR and TNR differences between the general model and (a) manifestation-specific models and (b) motion-specific models. The average ***±*** standard deviation is annotated. Manifestation-specific models improve the overall TPR by **24.95%±9.80%** and TNR by **18.29%±8.71%**, while motion-specific models decrease the overall TPR by **1.89%±8.74%** and TNR by **5.17%±10.76%**

**Fig. 6.**
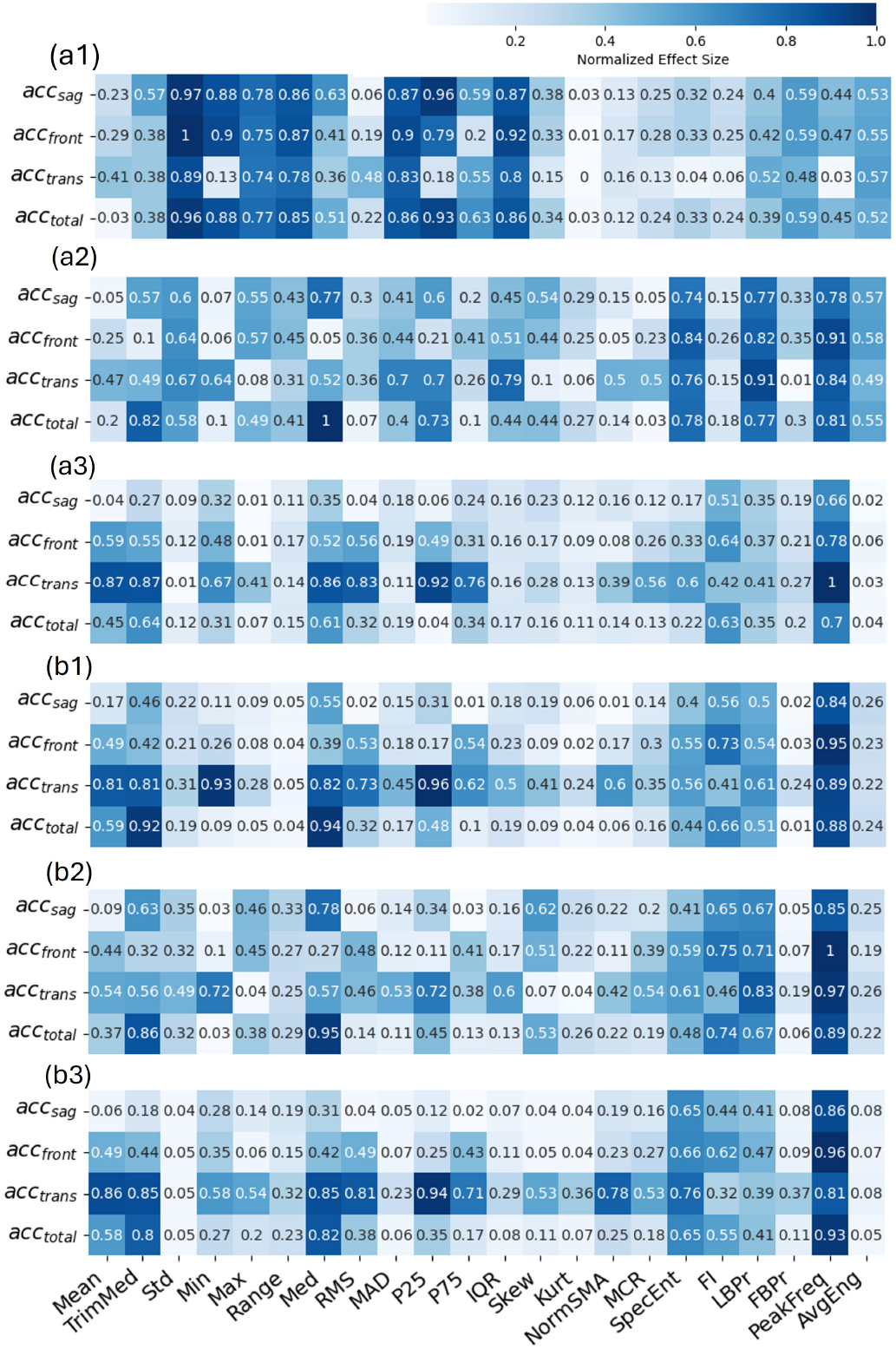
Average feature masks obtained with all non-FOG and (a) manifestation-specific subtypes ((a1) Akinesia, (a2) Shuffling, (a3) Trembling) and (b) motion-specific subtypes ((b1) Gait-initiation, (b2) Turning, (b3) Walking). It displays similar trends as subtype-group feature masks, especially that manifestation-specific feature masks (a) differ notably from each other and the general mask, whereas motion-specific feature masks (b) resemble each other and the general mask.

#### 1) ART ANOVA Test

Based on *p* values of Table II, the TPRs and TNRs on manifestation test sets are significantly influenced by the model and subtype test set, which justifies further comparisons of models within each subtype test set. The TPRs and TNRs on motion test sets are however only influenced by the test set.

#### 2) Manifestation-Specific Model Performances

For model performances in manifestation groups, with TNRs (specificity) shown in Fig. 4 (a) and TPRs (sensitivity) in (b), the general model has high specificity (i.e., correct non-FOG detection) for shuffling, moderate specificity for akinesia, and low specificity for trembling (typically below 50%). Akinesia and trembling models significantly improve non-FOG detection in the corresponding subtype groups, particularly trembling. This comes with a drop in trembling sensitivity, though the resulting sensitivity average remains around 82.93%, while no significant reduction is observed in akinesia sensitivity. In terms of sensitivity, the general model performs near-perfectly on akinesia and trembling but shows a marked weakness in shuffling. The shuffling model effectively resolves this issue, enhancing the mean TPR from 53.38% ± 7.18% to 82.06% ± 3.08%, at the cost of a moderate drop in the same-group TNR. To further quantify the average improvements by the subtype models, Fig. 5 (a) indicates the TPR and TNR differences between subtype models and general model, such that positive bars indicate an improvement. This figure illustrates that above-zero bars outweigh the below-zero bars in all subtype groups. To evaluate the overall performance changes by subtype models, the last column aggregates the TPR and TNR differences across all subtype test sets, treating each as equally important. It shows a noticeable net improvement on both TPR and TNR by 18.29%±8.71% and 24.95%±9.80%, respectively.

#### 3) Motion-Specific Model Performances

For model performances in motion groups, with TNRs illustrated in Fig. 4 (c) and TPRs in (d), the general model has a moderate median TPR and TNR (above 75%) on all test sets, while being more variant for walking FOG. Motion subtype models show no significant performance improvements compared to the general model except a small enhancement in sensitivity during turning, with a similar amount of specificity decline. According to Fig. 5 (b), there are more degradations than improvements by these subtype models while the extent of changes are small. The aggregated performance shifts by the motion models show a net degradation in TPR by 1.89%*±*8.74% and TNR by 5.17%±10.76%.

## IV. Discussion

### A. Feature Masks Reflect Dominant Subtype Group(s)

Subtype-specific feature masks reflect the traits of corresponding subtype groups, effectively representing targeted detection strategies. Akinesia feature mask aims to distinguish completely still freezing from mainly quite standing (akinesia-like non-FOG). Both cases are characterized by low, uniform frequency components, making FD features less helpful. Therefore, the attentions are shifted to TD features, such as MAD and IQR. Shuffling feature mask differentiates short, dragging steps from normal walking and turning (shuffling-like non-FOG), a challenge given their similar underlying gait patterns. Consequently, more TD and FD features are required to encode the subtle differences. For instance, greater reliance on FD features such as MCR helps capture gait rhythm differences, while increased attention to TD features such as range and Std helps describe gait speed and variability. Trembling feature mask separates tremor from sensor noise or jolting normal motion (trembling-like non-FOG), which also shares similar dominant FD characteristics and thus leads to increased attention to TD. Among the added TD features, those in transverse-plane are favored, as they capture directional changes and movement magnitude, both key for distinguishing true movement within noisy non-FOG windows.

The three motion feature masks exhibit similar FD-dominated feature preferences, with minor difference in walking case due to the addition of MCR, which may capture gait rhythm and tremor-related oscillations associated with walking FOG.

The general feature mask reflects its dominant subtype group(s), as quantitatively supported by correlation analysis, suggesting a potential bias towards the dominant subtype. It shows the highest Spearman and (or) Pearson correlations with the shuffling and trembling feature masks among all manifestation-specific feature masks. In the dataset, the non-FOG instances of the shuffling group, i.e., walking and turning, account for majority of normal motion, while trembling is the dominant manifestation subtype. Similarly, among the motion-specific feature masks, the general feature mask is most correlated with the turning feature mask, consistent with turning being the most prevalent motion context in the dataset.

### B. Manifestation Models Result in Improved Generalizability

As highlighted in Sec. IV-A, the general model biases toward the dominant FOG manifestation (trembling FOG) and the dominant non-FOG patterns (shuffling-like non-FOG), while overlooking less frequent non-FOG scenarios and underperforming on shuffling FOG. This bias arises because shuffling-like non-FOG not only dominates the non-FOG class but constitutes the majority of the overall dataset. Resolving this, manifestation models significantly improve detection sensitivity in shuffling and reduces false alarms in akinesia-like and trembling-like non-FOGs. As is common in binary classifiers, slight adjustment to the decision boundary can lead to improvement in one class at the expense of degradation in the other. Thus, the observed improvements are accompanied by a decrease in specificity in shuffling subtype and reduction in sensitivity in trembling subtype. However, the overall improvements outweigh these degradations. Consequently, the overall average TPR and TNR improve, indicating enhanced generalizability across the FOG subtypes and their corresponding non-FOG subsets. Notably, the improvements typically arise where the general model performs close to chance level (median TPR/TNR *<* 0.6), while degradations still retain acceptable performance (median TPR/TNR *>* 0.7), thereby justifying the trade-off.

In contrast, motion models only show statistically better sensitivity in turning FOG, albeit being small and leading to a similar specificity reduction for the same group. Overall performance shifts also show no improvements compared to the general model. This is expected since those subtype models and the general model share the same detection strategy as indicated by feature masks, making improvements via motion-specific tailoring unlikely.

### C. The Manifestation Composition of a Dataset Drives the Detection Strategy

Tailored training is effective only if the targeted subset is sufficiently distinct from the general data. Since the tailored training is effective with manifestation models but not motion models, it suggests that waist acceleration differ remarkably across manifestation groups but remain similar across motion groups. Thus, manifestation groups can act as building blocks of a detection strategy: a specific manifestation composition (i.e. proportion of samples in each manifestation group) of a dataset leads to a specific desired detection strategy that maximizes the overall accuracy.

Overall accuracy naturally favors dominant manifestation groups without accounting for subtype importance. This explains why a general model, optimized for the dataset’s overall manifestation distribution, tends to favor dominant manifestation(s). Motion-specific groups exhibit similar manifestation composition to the full dataset, as indicated by the shared detection strategy between the motion models and the general model. Since the general model is already tuned to this distribution, it performs reasonably well across all motion test sets, making motion-specific tailoring generally unnecessary. However, this does not rule out the possibility of further improving generalization across motion-specific groups, especially in presence of atypical events with distinct manifestation compositions. In such cases, a motion group can be further split to represent the underrepresented manifestation composition. For instance, motivated by the high variability of walking FOG sensitivity by both the general and walking model, researchers may further scrutinize walking group. Given the low sensitivity of the general model to shuffling, those walking test folds with low walking sensitivity likely contain more shuffling-dominated walking FOG than the general case. Being underrepresented, that atypical composition fails to influence the detection strategy and thus remains undetected. To address this, training a model specifically tailored to such less frequent manifestation profiles may enhance detection performance.

### D. Limitations and Future Directions

The utilized dataset has a minimal sensor setting of only one waist-placed accelerometer. Additional FOG-modulated bio-signals collected by other sensors such as skin conductance reflecting anxiety state [40] and foot plantar pressure reflecting both spatial and temporal gait patterns [41] may provide additional insights into the subtype-specific features and differences in FOG detection strategies. Collecting inertial data (acceleration and angular velocity) from other body segments, for instance, ankles, may help further improve deep learning model accuracy [29]. Additionally, the manifestation-specific subtype labels are from our clustering-based split approach instead of expert annotations. Expert labels may improve the quality of dataset and lead to more rigorous manifestation categorization. Moreover, this study focuses on demonstrating the need for subtype-specific strategies rather than developing an optimal subtype model fusion. XGBoost is a practical proxy for motion-specific model activation, though its suboptimal test classification accuracy leaves room for refinement in future fusion approaches. Future work may also apply the same verification method to the FOG prediction problem, enabling proactive interventions.

## V. Conclusion

This study demonstrates that subtype-specific strategies can significantly enhance FOG detection, particularly when tailored to manifestation-based subtypes. Through feature mask analysis, we revealed that different FOG manifestations (akinetic, shuffling, trembling) require distinct detection strategies, while motion-based subtypes (gait initiation, walking and turning) share more homogeneous strategies. To leverage this insight, we then developed feature-mask-based CNNs that explicitly embed subtype-specific strategies these derived strategies derived from interpretable feature masks. Models trained on manifestation-specific subtypes significantly outperformed a general feature-masked-based CNN model, indicating improved generalizability across subtypes. In contrast, motion-specific models do not yield such improvement, as they share similar detection strategies with the general model. This supports our conclusion that FOG detection strategies are driven more by manifestation composition than motion context. Future work may explore more comprehensive sensor configurations and clinically informed subtype definitions and data split to further refine subtype-sensitive detection models.

