## Supplementary Material for "Feature-Mask-Based Strategies for Subtype-Specific Freezing of Gait Detection using CNNs" (from another version of this preprint: version 1, 2025-05-29)

Thea Yu, Kaylena Ehgoetz Martens, Arash Arami

The supplementary material contains samples of waist-worn accelerometer signals of manifestation-specific subtypes and their same-cluster non-FOG subsets (Fig. S1), motion-specific subtypes and their same-cluster non-FOG subsets (Fig. S2). It also depicts the average feature masks obtained with each FOG subtype and all Non-FOG data (Fig. S3).

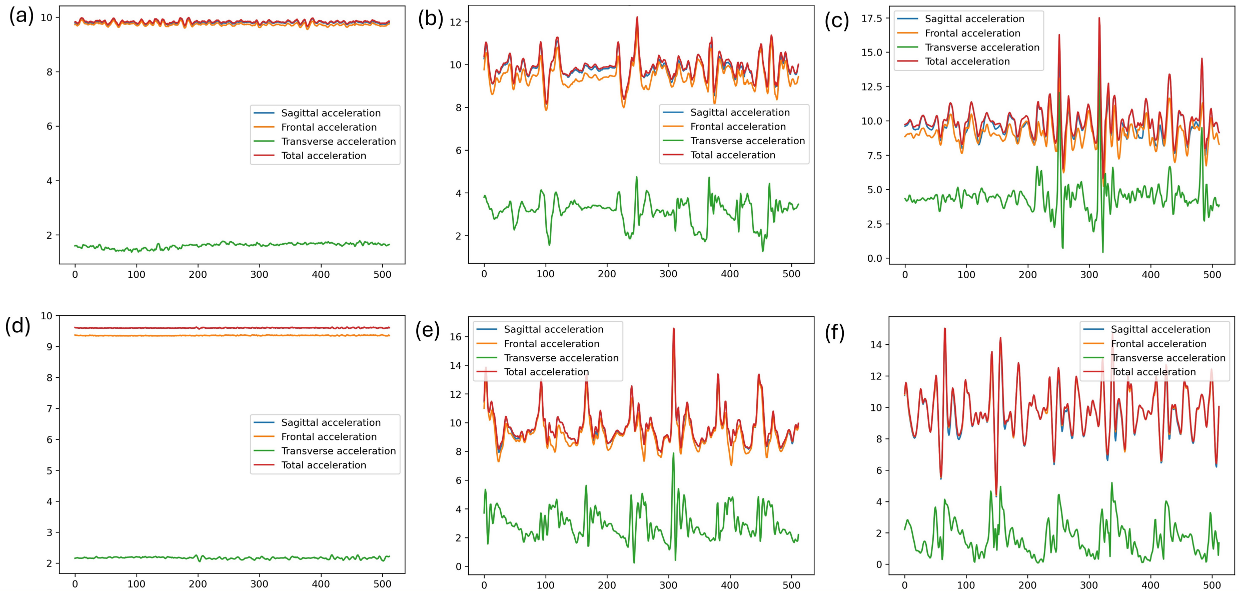


Figure S1: Example Sample of (a) Akinesia FOG (b) Shuffling FOG (c) Trembling FOG (d) Akinesia-like Non-FOG (e) Shuffling-like Non-FOG (f) Trembling-like Non-FOG

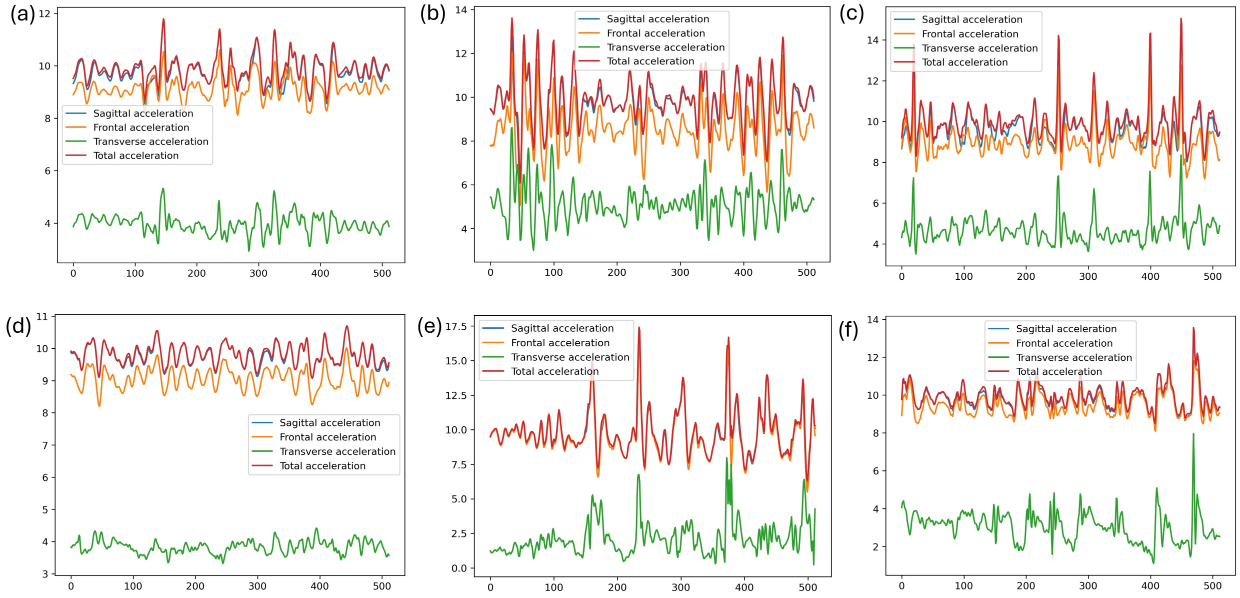


Figure S2: Example Sample of (a) Gait-initiation FOG (b) Turning FOG (c) Walking FOG (d) Gait-initiation-FOG-like Non-FOG (e) Turning-FOG-like Non-FOG (f) Walking-FOG-like Non-FOG

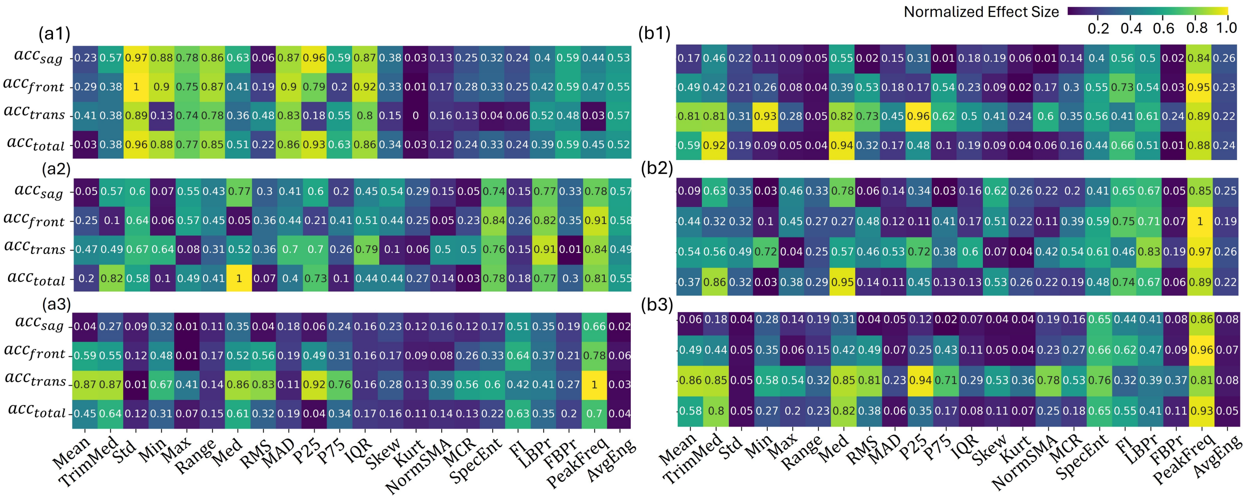


Figure S3: Average Feature Mask Obtained with FOG Subtype and All Non-FOG for (a) manifestation-specific subtypes ((a1) Akinesia, (a2) Shuffling, (a3) Trembling) and for (b) motion-specific subtypes ((b1) Gait-initiation, (b2) Turning, (b3) Walking)
